# Communities of tropical soil fungi differ between burned and unburned forest, with corresponding changes in plant community composition, litter and soil chemistry

**DOI:** 10.1101/2021.08.22.457293

**Authors:** Jed Calvert, Alistair McTaggart, Lília C. Carvalhais, André Drenth, Roger Shivas

**Author notes:** Corresponding author: Centre for Horticultural Science, Queensland Alliance for Agriculture and Food Innovation, The University of Queensland, Ecosciences Precinct, 41 Boggo Rd Dutton Park, Queensland 4102, Australia. E-mail address (J. Calvert).

## Abstract

Fire is predicted to be more severe and frequent in forests of the Australian Monsoon Tropics over the coming decades. The way in which groups of ecologically important soil fungi respond to disturbance caused by fire has not been studied in Australian tropical forest ecosystems. Ectomycorrhizal (EM) fungi are important tree symbionts and saprotrophic fungi drive soil nutrient cycles. We analysed both publicly-available environmental DNA sequence data as well as soil chemistry data to test a hypothesis that burned areas in a contiguous tropical forest have different community composition and diversity of EM and saprotrophic soil fungi relative to nearby unburned sites. We tested this hypothesis by measuring community-level taxonomic composition, fungal diversity, species richness and evenness. We determined whether changes in fungal communities were associated with fire-altered soil chemical/physical properties, vegetation types, or the direct effect of fire. Soil fungi differed in abundance and community phylogenetic structure between forest sites that had experienced fire, and sites dominated by unburned forest. EM community composition differed between unburned and burned sites, which had more mycorrhizal hosts including *Corymbia intermedia, Acacia flavescens* and *Acacia midgleyi*. Lower diversity of saprotrophic fungi was correlated with lower soil nutrient levels and different litter composition at burned sites. Pyrophilic, truffle-like EM fungi that rely on mycophagous mammals for dispersal were abundant at recently burned sites. We conclude that EM fungi show different patterns of diversity in burned tropical forest, likely driven by changing plant communities, whereas differences in saprotrophic fungal communities of burned sites may be driven by by reduced soil nutrient levels and altered litter composition.

**Graphical abstract:** 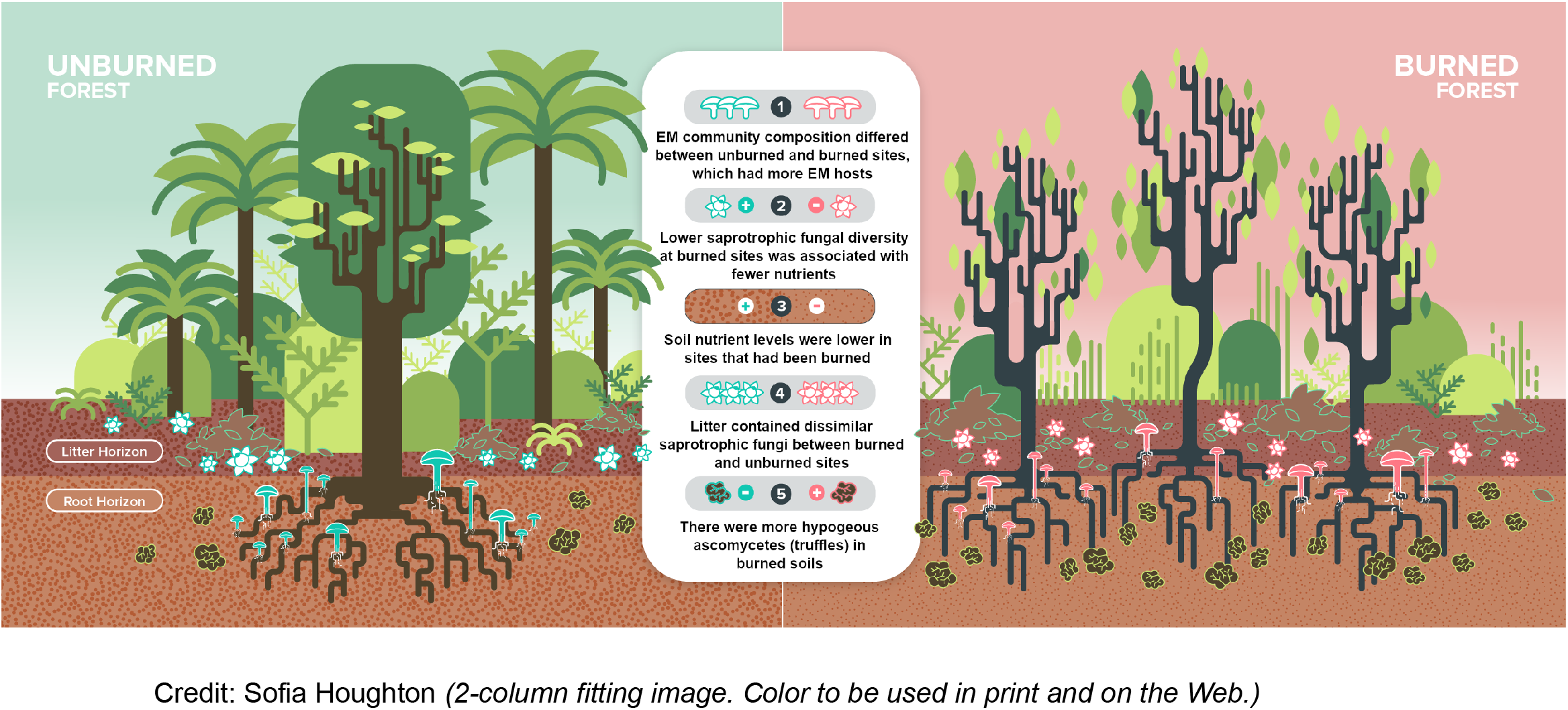

## 1. Introduction

Fire is a major driver of disturbance in tropical forests (Silvério et al., 2019), including the Australian Monsoon Tropics (AMT), which is the most fire-prone region in Australia (Bowman et al., 2010). In this region, fire frequently intrudes from savanna into notophyll plant communities and initiates a process of post-fire seral development (Bowman, 2009; Cole et al., 2014). Fire is a primary determinant of plant distribution, which in turn can influence the structure of microbial communities (Ettema and Wardle, 2002; Ondei et al., 2016; Sarmiento et al., 2017). An increase in fire frequency, severity and duration in the AMT is likely over the next decades due to anthropogenic climate change (Hubnerova et al., 2020).

Soils contain some of the most complex and understudied ecosystems in terrestrial biomes, providing habitat for an estimated 25 % of described species (Decaëns et al., 2010). Most of the terrestrial carbon on Earth is in soils (Crowther et al., 2016), and they have been designated a ‘third biotic frontier’ after deep-sea benthic regions and tropical rainforest canopies (Hågvar, 1998). Healthy soil ecosystems are undergirded by diverse communities of microorganisms dominated by fungi, bacteria, archaea and other eukaryotes, the taxonomy and function of which are largely unknown (Baldrian, 2019). Together, the microorganisms of this ‘living terrestrial skin’ drive global biogeochemical cycles and power terrestrial ecosystems (Tecon and Or, 2017).

Specific functional guilds of microorganisms respond differently to fire according to their trophic modes. The resilience of tropical ectomycorrhizal (EM) fungi after fire has been attributed to their ability to draw nutrients from plant roots (Alem et al., 2020). Fires impact soil microorganisms through changes in soil pH, water holding capacity, and availability of organic carbon, nitrogen and phosphorus (Pellegrini et al., 2019; Singh, 1994; Verma and Jayakumar, 2018). Soil enzyme activity, which reflects microbial metabolism in soil communities, also decreases immediately following fires, especially at shallow soil horizons (Certini et al., 2021). How different functional guilds of fungi respond to fire in an Australian tropical soil ecosystem has not been studied.

Saprotrophic and EM fungi are two functional guilds of fungi in tropical forests that break down soil organic matter (SOM), with saprotrophs degrading SOM primarily as a source of carbon, and EM fungi using it as a source of nutrients (Fernandez and Kennedy, 2016; Lindahl and Tunlid, 2015). Gadgil and Gadgil (1975, 1971) proposed that competition and inhibition between saprotrophic and EM fungi suppress the decomposition of organic matter and increase the accumulation of organic carbon. Studies in northern-hemisphere coniferous ecosystems based on post-fire observations of macrofungal sporocarps have reported lower EM diversity (Owen et al., 2019) and higher saprotroph diversity (Salo et al., 2019). The abundance of tropical arbuscular mycorrhizal fungi is slightly reduced by fires based on counts of spores and propagules (Aguilar-Fernández et al., 2009; Raman and Nagarajan, 1996). Studies in coniferous ecosystems have reported dominance of saprotrophic fungi in unburned areas, which decreased in diversity after fire (Semenova-Nelsen et al., 2019), as well as long-term reduction of both ectomycorrhizal and saprotrophic diversity after fire (Martín-Pinto et al., 2006; Pulido-Chavez et al., 2021). Some of this reduction in fungal diversity may be a result of large increases in abundance of some Ascomycete groups (Reazin et al., 2016), and several studies have estimated that more than a decade may be required for soil ecosystems to return to a state similar to unburned soils (Dooley and Treseder, 2012; Oliver et al., 2015). In Australian tropical forests, it remains unclear whether communities of saprotrophic and EM fungi show different patterns of diversity between burned and unburned sites.

We used data from culture-independent high-throughput sequencing of soils, provided by the Biomes of Australian Soil Environments (BASE) soil microbial diversity database, to test a hypothesis that burned areas in a contiguous tropical forest have different community composition and diversity of EM and saprotrophic soil fungi relative to nearby unburned sites. BASE has mapped Australian soil microbial diversity using culture-independent high-throughput DNA sequencing (Bissett et al., 2016). We measured community-level taxonomic composition, diversity, species richness and evenness. We tested whether observed changes were associated with the burn status (burned/unburned) of a site, the recency of a fire, and whether these changes were correlated with altered soil chemical/physical properties or vegetation type between burned and unburned sites. We hypothesised that changes would be correlated with altered soil chemical/physical properties or vegetation type. Understanding the response of soil microbe communities to fire in the AMT may provide management options for the protection of ecosystems under a changing climate.

## 2. Materials and methods

### 2.1. Study site

The Iron Range on Cape York Peninsula, Far North Queensland, is a hilly coastal region of the Australian Monsoon Tropics (AMT) dominated by tropical rainforest and notophyll vine forest (Neldner and Clarkson, 1995; Webb, 1959). Sample sites for this study (−12.779910331934488, 143.3128161670805) were within a radius of 750 m and were selected to represent a spectrum of seral stages, from unburned to recently burned (*Figure 1, Table 1*). Site elevation ranged from 40 m to 94 m above sea level, with a mixture of moderately deep gradational yellow soils formed on hillslopes of adamellite or granite, and deep gradational brown structured soils derived from schist, phyllite, quartzite and gneiss (QSpatial, 2020).

**Figure 1.**
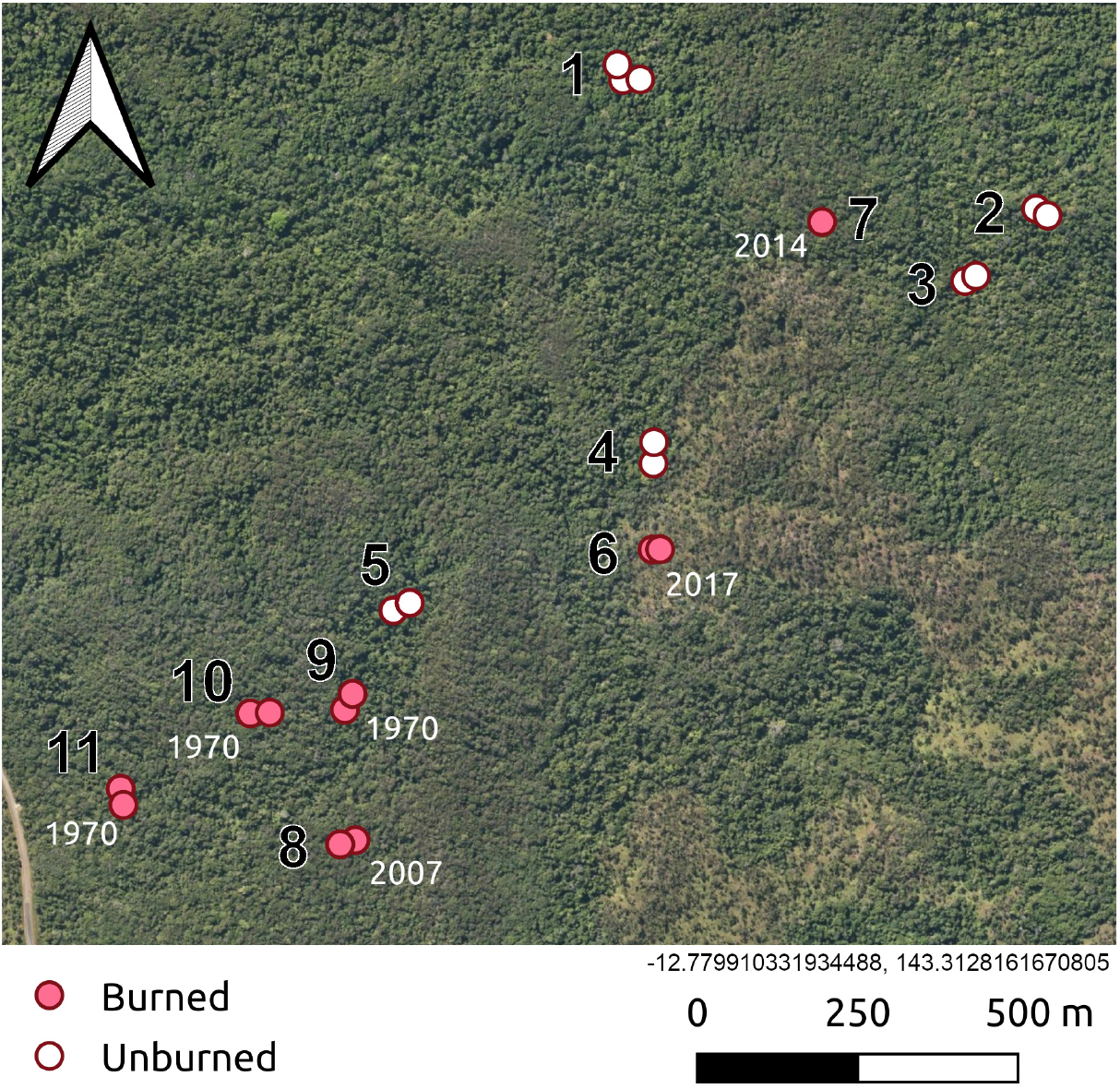
Soil sampling locations. Study sites numbered in black type, with year of most recent fire at burned sites marked in white type. Each point represents 2 samples, one from litter (0–10 cm) and mineral soil (20–30 cm). (*Single-column fitting image. Color to be used in print and on the Web*.)

**Table 1.** Study sites, fire history and floristic composition. Plant species listed are arbuscular mycorrhizal (AM) unless marked with an * (EM-AM, ectomycorrhizal & AM) or ^ (NM/AM, non-mycorrhizal or AM). (*2-column fitting table*.)

| Site | Most recent fire | Vegetation | Dominant trees | Dominant grasses |
| --- | --- | --- | --- | --- |
| 1 | Unburned | Evergreen notophyll vine forest | *Aleurites moluccanus*, *Neonauclea glabra*, *Canarium australium* var. *aust.*, *Archidendron hirsutum*, *Beilschmiedia obtusifolia* |  |
| 2 | Unburned | Semi-deciduous mesophyll/notophyll vine forest | *Cordia dichotoma*, *Tetrameles nudiflora*, *Berrya javanica*, *Paraserianthes toona*, *Mimusops elengi*<sup>^</sup> |  |
| 3 | Unburned | Semi-deciduous mesophyll/notophyll vine forest | *Tetrameles nudiflora*, *Cordia dichotoma*, *Canarium australium* var. *aust.*, *Lagerstroemia archeriana*, *Vitex helogiton* |  |
| 4 | Unburned | Evergreen notophyll vine forest | *Terminalia complanata*, *Palaquium galactoxylum*, *Garcinia dulcis*, *Syzygium pseudofastigiatum*, *Endiandra impressicosta* |  |
| 5 | Unburned | Semi-deciduous mesophyll/notophyll vine forest | *Tetrameles nudiflora*, *Blepharocarya involucrigera*, *Palaquium galactoxylum*, *Alstonia scholaris*, *Aleurites moluccanus* |  |
| 6 | 2017 | Woodland/grassland | *Corymbia intermedia*<sup>*</sup>, *Lophostemon suaveolens*<sup>*</sup>, *Acacia flavescens*<sup>*</sup>, *Planchonia careya*, *Grevillea parallela* | *Heteropogon contortus*, *Imperata cylindrica*, *Heteropogon triticeus*, *Entolasia stricta*, *Gahnia aspera*<sup>^</sup> |
| 7 | 1995, one in 2014 | Monodominant post-fire | *Dodonaea viscosa* |  |
| 8 | 1970, one in 2006 | Regenerating closed canopy rainforest | *Dillenia alata*, *Buchanania arborescens*, *Guioa acutifolia*, *Blepharocarya involucrigera*, *Litsea breviumbellata* | *Cyrtococcum oxyphyllum*, *Ancistrachne uncinulata*, *Entolasia stricta* |
| 9 | 1970 | Edge of intact and regenerating rainforest | *Nauclea orientalis*, *Buchanania arborescens*, *Vitex helogiton*, *Alphitonia* sp., *Syzygium forte* subsp. *forte* |  |
| 10 | 1970 | Regenerating closed canopy rainforest | *Phyllanthus praelongipes*, *Mallotus resinusus*, *Mallotus polyadenos*, *Rinorea bengalensis*, *Buchanania arborescens* |  |
| 11 | 1970 | Regenerating closed canopy rainforest | *Atractocarpus sessilis*<sup>^</sup>, *Buchanania arborescens*, *Acacia midgleyi*<sup>*</sup>, *Mallotus philippensis*, *Larsenaikia ochreatea* |  |

### 2.2. Sampling BASE data

We analysed 44 fungal amplicon community profiles from 5 unburned and 6 burned sites from the Biomes of Australian Soil Environments microbial diversity database (BASE), which were sampled and sequenced according to Bissett et al. (2016). Between 2 and 6 soil samples were taken from each site (see *Figure 1*). The BASE project took soil samples of 1 kg from the litter (0–10 cm) and mineral soil (20–30) in February 2017. Soil chemical/physical properties including ammonium (NH_4_), nitrate (NO_3_), phosphorus (P), potassium (K), organic carbon (C), calcium (Ca) and pH were analysed and DNA was extracted from samples as per protocols of the Earth Microbiome Project (*http://www.earthmicrobiome.org/emp-standard-protocols/dna-extraction-protocol/*). The ITS region of fungal ribosomal DNA was amplified with the primers ITS1F and ITS4 (Gardes and Bruns, 1993; White et al., 1990) and sequenced with 300 bp paired-end chemistry on an Illumina MiSeq, retaining the only the ITS1 region by using only forward reads (Bissett et al., 2016). Plant mycorrhizal traits were established using the FungalRoot global online database of plant mycorrhizal associations’ *Recommended mycorrhizal status for plant genera* (Soudzilovskaia et al., 2020).

### 2.3. Processing of sequence data

ITS1 reads were identified and extracted with ITSx v1.1.3 (Bengtsson-Palme et al., 2013). Quality filtering and construction of operational taxonomic unit (OTU) tables were performed in QIIME2 v2020.11 (Bolyen et al., 2018) with the `dada2 denoise-single, phylogeny align-to-tree-mafft-fasttree, diversity core-metrics-phylogenetic` and `featureclassifier classify-sklearn` functions. OTUs were generated from sequences with 97 % similarity, and taxonomy was assigned using the UNITE v8.2 fungal database (Abarenkov et al., 2010). Fungal community diversity was calculated from the ITS dataset rarefied to 5,000 sequences per sample, based on rarefaction curves of Shannon’s diversity index.

### 2.4. Statistical analyses

Soil chemistry data for each site were analysed to establish whether nutrient content was correlated between samples exposed to fire at different time points and fungal community structure. A distance matrix of nutrient profiles for each site was constructed in R v3.6.3 (R Core Team, 2020) based on Bray-Curtis dissimilarities (Bray and Curtis, 1957) with the function `vegdist` in Vegan v2.5-6 (Oksanen et al., 2020) and visualized with non-metric multidimensional scaling (NMDS) (function `metaMDS`). Soil chemical/physical properties were analysed for NH_4_, NO_3_, P, K, C, Ca and pH. To establish whether fire history and other factors structured soil fungal communities, we constructed distance matrices from OTU tables based on unweighted UniFrac (Lozupone and Knight, 2005), which measures OTUs in terms of their phylogenetic relatedness and presence or absence between samples. We built PERMANOVA (`adonis`) forward models in R to assess variance between categorical variables related to soil chemical/physical properties and determine the significance and hierarchy of influence for sample depth, burn status (burned/unburned), vegetation type (shrubland, grassy woodland, regenerated closed canopy forest, semi-deciduous notophyll forest, wet rainforest), year of most recent fire and year of cessation of frequent fires. We visualized Bray-Curtis distances based on soil chemical/physical properties at each sample site with NMDS (`metaMDS`) in Vegan to assess differences between fungal communities in terms of soil chemical/physical properties.

Sequences that represented ectomycorrhizal and saprotrophic fungi were identified with the FUNGuild v1.1 (Nguyen et al., 2016) Python script on an ITS OTU table rarefied to 5,000 sequences and with singletons removed. Only ‘probable’ and ‘highly probable’ assignments were retained. To detect linear correlations between sample alpha diversity and soil chemical/physical properties, we generated Shannon’s diversity (entropy) values (Shannon, 1948) for all samples in QIIME2 (`qiime diversity alpha`) and Pearson’s correlation coefficient, which measures the strength of a linear relationship between two variables, with rcorr in the R package Hmiscv4.4-2 (Harrell, 2021). We used redundancy analysis (RDA) in R with the package GGORD (Beck, 2017) to extract and summarise the variation in response variables (sample fungal community composition, individual taxa) and explanatory variables (burn status, soil chemical/physical properties) based on Hellinger-transformed OTU tables to give lower weights to rare taxa. Krona v2.7.1 (Ondov et al., 2011) was used to visualise the proportional taxonomic composition of fungal communities.

## 3. Results

### 3.1. Sequence data

After DADA2 quality filtering in QIIME2, we retained 722,732 ITS sequences from 42 samples, which clustered into a total of 6960 fungal OTUs, from which we identified 165 EM and 654 saprotrophic taxa.

### 3.2. Soil chemistry

PERMANOVA indicated that sample depth (P=0.001) and burn status, i.e. whether or not a site had been burned (P = 0.012) had the most influence on fungal communities (*Table 2*). Interactions between variables were associated with differences in soil chemistry. Depth interacted with burn status (P = 0.001), vegetation type (P = 0.001), year of cessation of frequent fires (P = 0.010) and year of last fire (P = 0.010). NMDS showed that samples clustered primarily with sample depth and burn status (*Figure 2*). The greatest variability in soil chemistry was in outlier samples from unburned mesophyll/notophyll rainforest.

**Figure 2.**
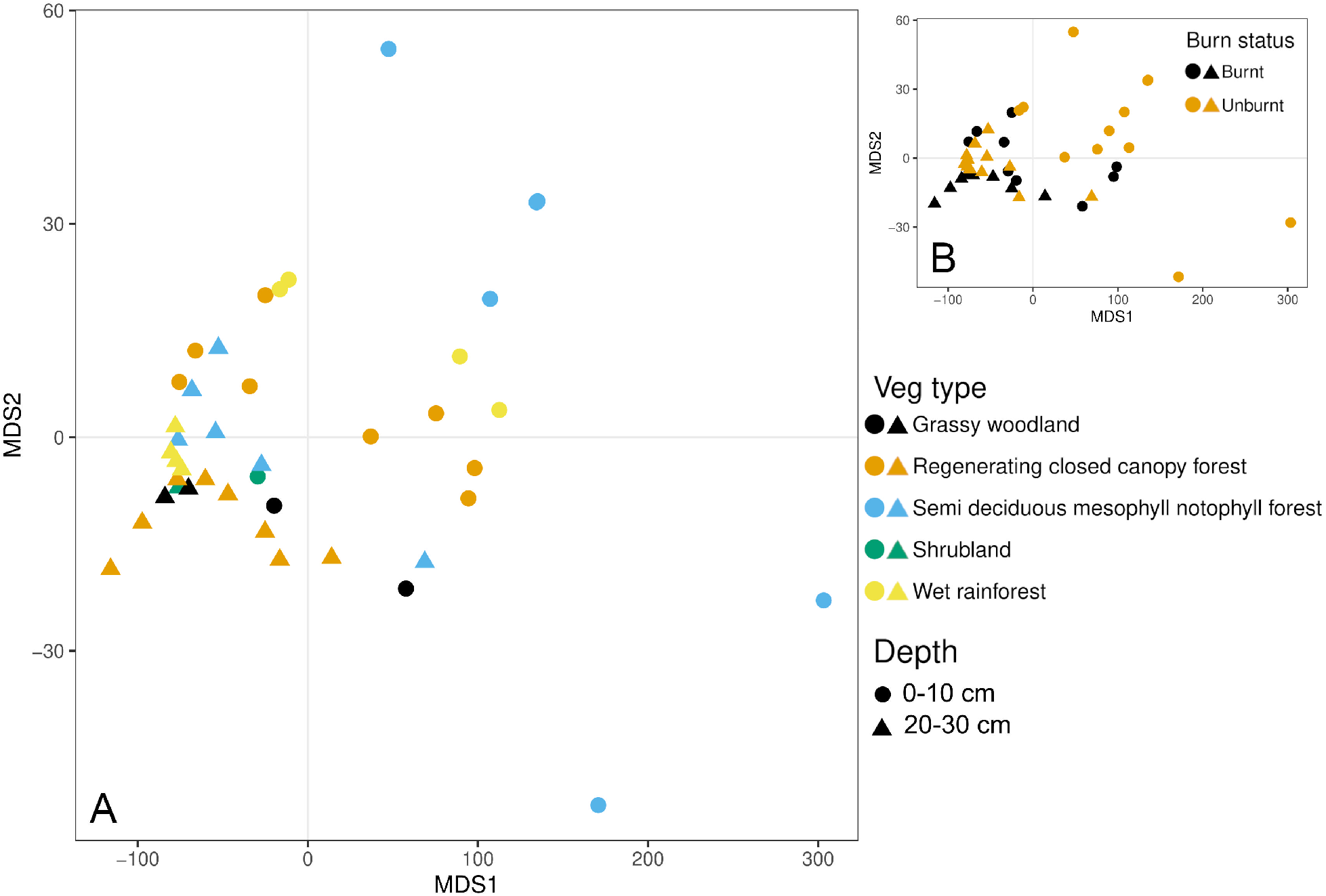
Non-metric multidimensional scaling showing differences between soil chemistry of samples from sites with different vegetation types and burn status (burned/unburned). Based on a Bray-Curtis distance matrix of nutrient variables for soils at each each site including ammonium (NH_4_), nitrate (NO_3_), phosphorus (P), potassium (K), organic carbon (C), pH and calcium (Ca). (*1.5-column fitting image. Color to be used in print and on the Web*.)

**Table 2.**
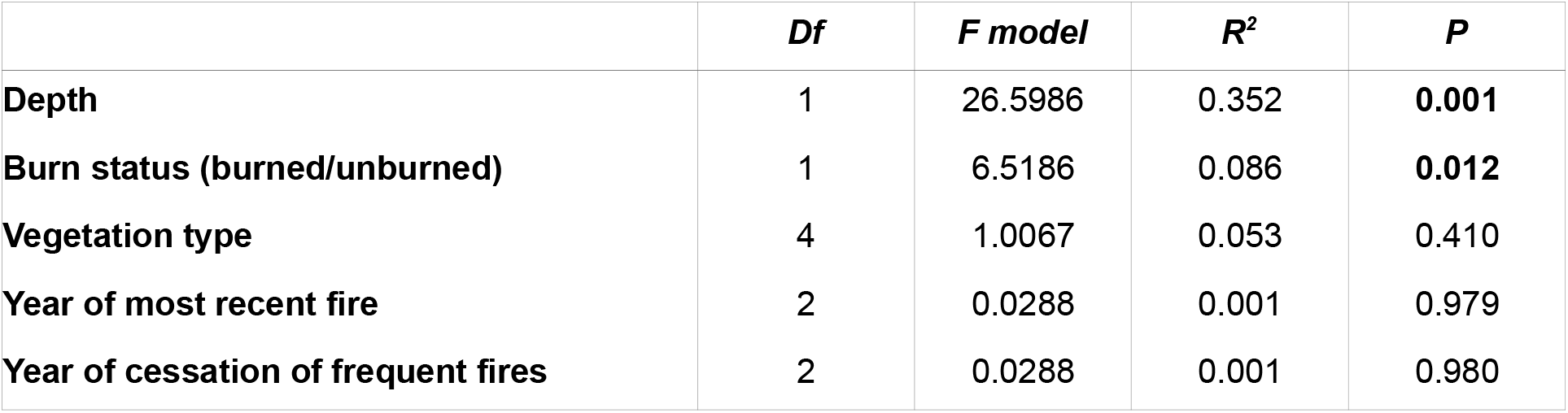
PERMANOVA of relative influence on soil chemical/physical properties of sampling depth, burn status (burned/unburned), vegetation type (shrubland, grassy woodland, regenerating closed canopy forest, semi-deciduous notophyll forest, wet rainforest), year of most recent fire and year of cessation of frequent fires. P values <0.05 indicated in bold type. (*Single-column fitting table*.)

### 3.3. Fungal community diversity and effects of soil nutrient levels

Shannon’s diversity index (entropy) was higher for saprotrophic fungi than EM fungi (*Figure 3*), particularly in the litter layer (0–10 cm) at unburned forest sites. Shannon’s diversity index of saprotrophic fungi correlated linearly with all measurements of soil physical/chemical properties. Diversity of EM communities correlated only with NH_4_ (*Table 3*). In general, NO_3_, P, Ca and pH were higher in unburned than in burned sites. Increased diversity of saprotrophic fungi correlated with levels of soil NH_4_, NO_3_, P, K and Ca, and there was a marginally significant correlation with pH. Lower diversity of EM communities at several unburned sites was associated with elevated soil nutrient levels relative to burned sites, although this trend was less evident for K levels (*Figure 4*). At unburned sites, saprotrophs were more diverse at 0–10 cm depth if levels of NO_3_, P, K and Ca were elevated. Saprotrophic diversity was more variable at 20–30 cm depth, where nutrient levels were lower.

**Figure 3.**
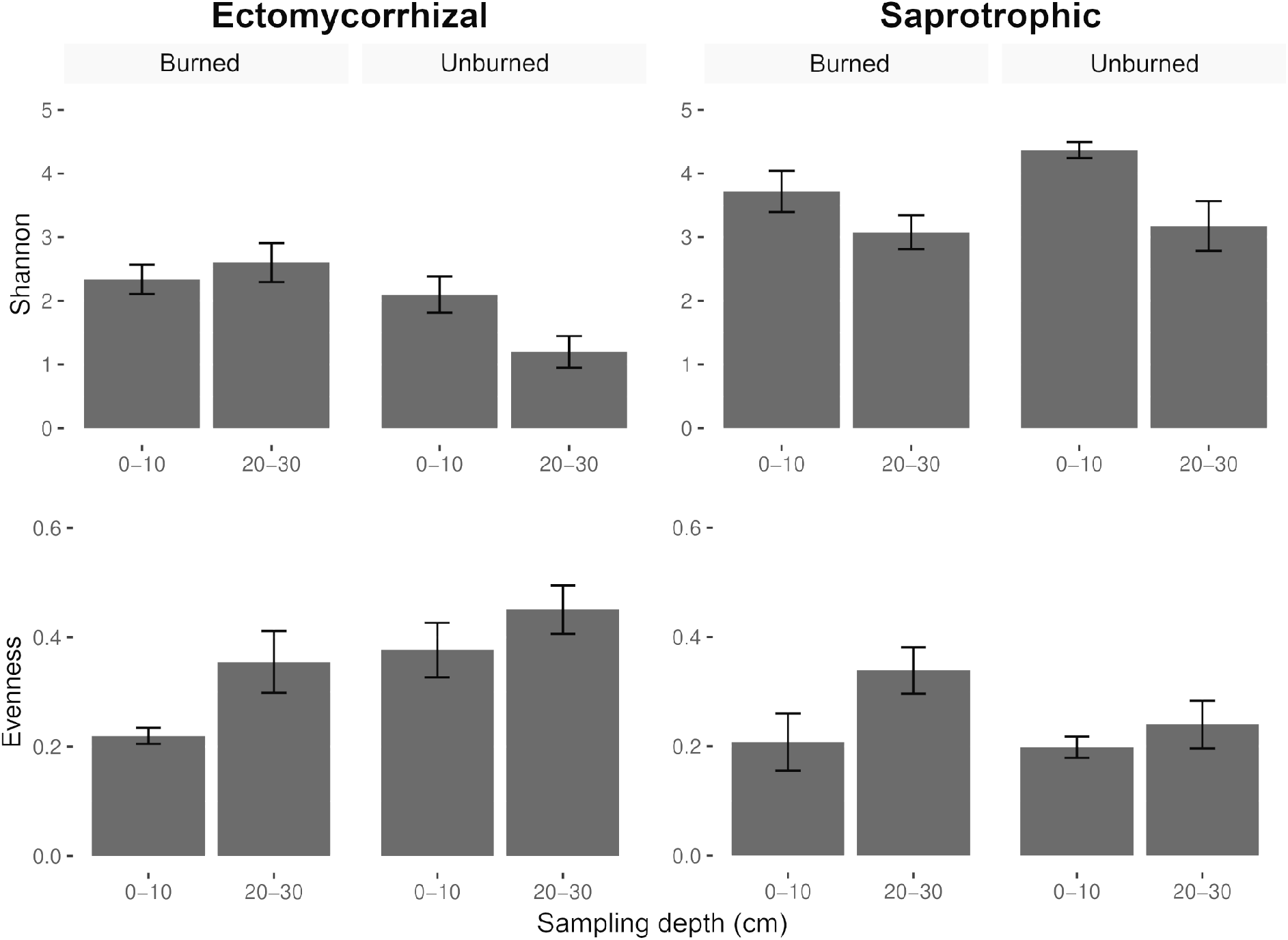
Average Shannon’s diversity values and Simpson’s evenness (similarity of abundance between species) for EM and saprotrophic soil fungi at burned and unburned sites and at two sampling depths. Bars indicate standard error. Shannon’s diversity was higher for saprotrophic fungi than EM fungi, particularly in the litter layer (0–10 cm) at unburned forest sites. EM communities had higher average Shannon’s diversity indices and lower evenness at burned compared to unburned sites (Figure 3). Saprotrophic communities were more diverse and less even at unburned sites. (*Single-column fitting image*.)

**Figure 4.**
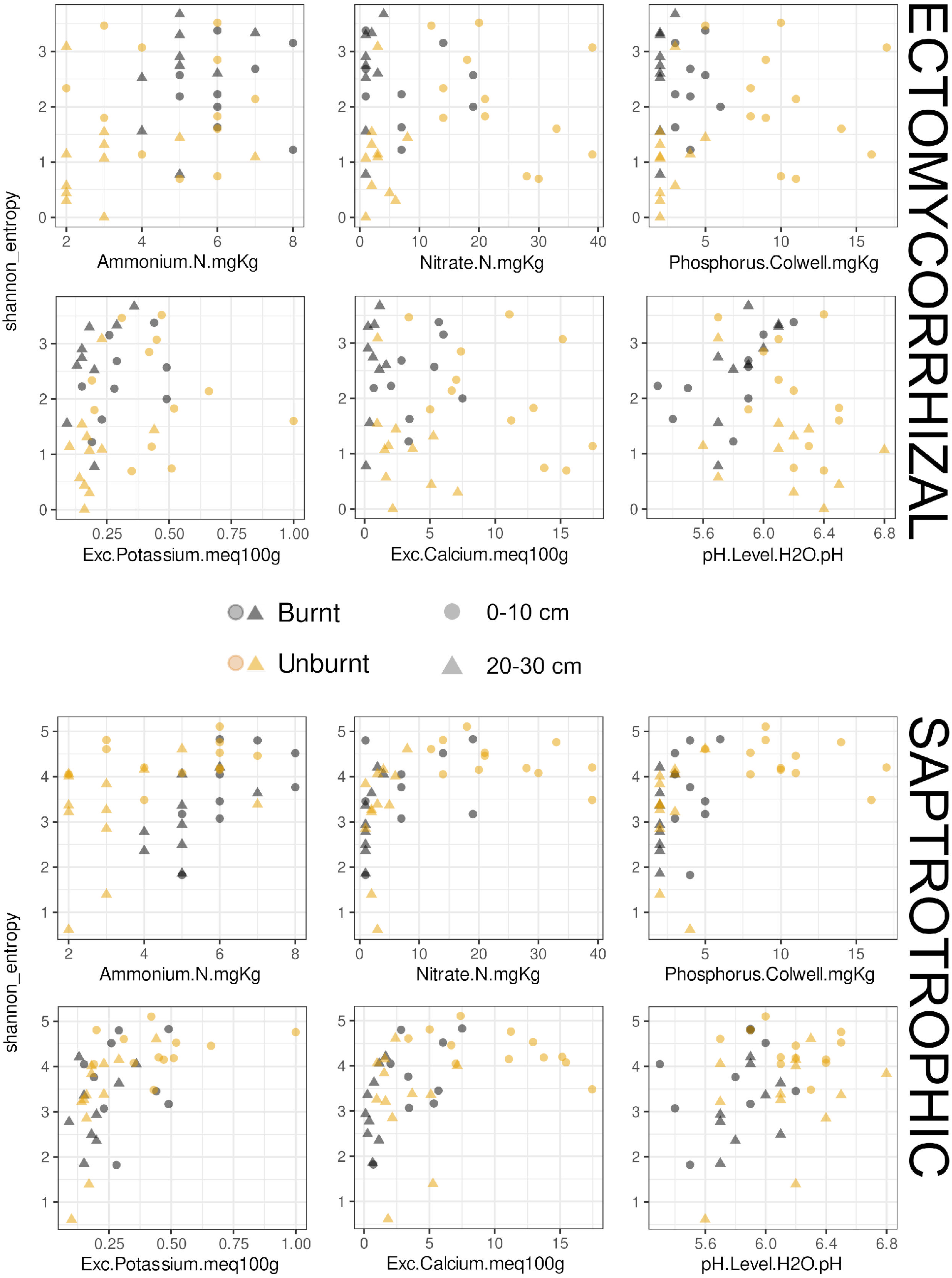
Shannon’s diversity (entropy) for ectomycorrhizal and saprotrophic soil fungi plotted against soil physical/chemical properties (levels of NO_3_, NH_4_, P, K, Ca & pH) at burned and unburned sites and at two sampling depths. Increased diversity of saprotrophic fungi correlated with levels of soil NH_4_, NO_3_, P, K and Ca, and there was a marginally significant correlation with pH. Lower diversity of EM communities at several unburned sites was associated with elevated soil nutrient levels relative to burned sites, although this trend was less evident for K levels. (*2-column fitting image. Color to be used in print and on the Web*.)

**Table 3.** Pearson’s correlation coefficient testing linear relationships between Shannon’s diversity (entropy) of soil fungal communities and soil chemical/physical properties. P values <0.05 indicated in bold type. (*Single-column fitting table*.)

|  | Ectomycorrhizal |  | Saprotrophic |  |
| --- | --- | --- | --- | --- |
|  | Pearson's correlation | P | Pearson's correlation | P |
| Ammonium | 0.330 | **0.035** | 0.326 | **0.035** |
| Nitrate | -0.018 | 0.913 | 0.464 | **0.002** |
| Phosphorus | 0.041 | 0.799 | 0.422 | **0.005** |
| Potassium | 0.186 | 0.243 | 0.511 | **0.001** |
| pH level | -0.236 | 0.138 | 0.299 | 0.055 |
| Calcium | -0.137 | 0.393 | 0.388 | **0.011** |

### 3.4. Fungal diversity and community structure and effects of historical burning

Whether or not a site had been burned was strongly correlated with changes in community structure of saprotrophic fungi at both sampling depths, and of EM fungi at 20–30 cm below the soil surface (*Table 4*). Sites 6 and 11, which were the only sites with EM host plants present, accounted for 20.1 % and 15.5 % of EM sequence reads, respectively. The number of years that had elapsed since the most recent fire had most influence on EM community structure 0–10 cm below the surface (P=0.005). Vegetation type was the second most important factor for EM fungi at 20–30 cm (P=0.011). Saprotrophic soil fungi at 0–10 cm depth were secondarily influenced by vegetation type (P=0.010). The influence of years since the most recent fire was marginally significant (P = 0.080) at 20–30 cm depth. An interaction was detected between vegetation type and the number of years since the most recent fire at 20–30 cm depth on the community structure of saprotrophic soil fungi (P=0.007).

**Table 4.** Results of PERMANOVA testing whether the burn status of a site (burned/unburned) and vegetation type (shrubland, grassy woodland, regenerating closed canopy forest, semi-deciduous notophyll forest, wet rainforest) were associated with changes in the community composition of ectomycorrhizal and saprotrophic soil fungi. Depth of sampling was a primary influence on community composition of EM (P=0.019) and saprotrophic (P=0.045) fungi, and datasets were split according to depth. All sites were included in the first analysis, as well as interactions between factors. Influence of years since most recent fire, years since cessation of frequent fires and interactions with vegetation type were subsequently analysed for burned sites only. P values <0.05 indicated in bold type. (*2-column fitting table*.)

|  | Depth (cm) | Factor | F model | R<sup>2</sup> | P |
| --- | --- | --- | --- | --- | --- |
| Ectomycorrhizal | 0–10 | **Years since most recent fire** | **1.93** | **0.22** | **0.005** |
|  |  | **Burn status** | **1.32** | **0.07** | **0.046** |
|  |  | Veg type | 0.81 | 0.17 | 0.635 |
|  |  | Burn status:Veg type | 0.73 | 0.20 | 0.797 |
|  |  | Years since cessation of frequent fires | 1.19 | 0.15 | 0.177 |
|  |  | Years since most recent fire:Veg type | 0.93 | 0.24 | 0.512 |
|  | 20–30 | **Burn status** | **1.37** | **0.07** | **0.007** |
|  |  | **Veg type** | **4.26** | **0.52** | **0.011** |
|  |  | **Burn status:Veg type** | **3.32** | **0.53** | **0.019** |
|  |  | Years since most recent fire | 1.01 | 0.13 | 0.552 |
|  |  | Years since cessation of frequent fires | 0.77 | 0.10 | 0.505 |
|  |  | **Years since cessation of frequent fires:Veg type** | **5.99** | **0.67** | **0.027** |
| Saprotrophic | 0–10 | **Burn status** | **1.67** | **0.08** | **0.004** |
|  |  | **Veg type** | **2.19** | **0.35** | **0.010** |
|  |  | **Burn status:Veg type** | **1.80** | **0.37** | **0.031** |
|  |  | Years since most recent fire | 1.01 | 0.13 | 0.549 |
|  |  | Years since cessation of frequent fires | 0.95 | 0.12 | 0.507 |
|  |  | Years since cessation of frequent fires:Veg type | 2.06 | 0.41 | 0.086 |
|  | 20–30 | **Burn status** | **1.39** | **0.07** | **0.046** |
|  |  | Veg type | 1.22 | 0.23 | 0.164 |
|  |  | Burn status:Veg type | 1.01 | 0.25 | 0.239 |
|  |  | Years since most recent fire | 3.28 | 0.32 | 0.080 |
|  |  | Years since cessation of frequent fires | 1.10 | 0.14 | 0.299 |
|  |  | **Years since most recent fire:Veg type** | **5.22** | **0.63** | **0.007** |

EM communities had higher average Shannon’s diversity indices and lower evenness at burned compared to unburned sites (*Figure 3*). Saprotrophic communities were more diverse and less even at unburned sites. Notably, the average Shannon’s diversity of EM fungi at unburned sites was lowest at deeper soil horizons (20-30 cm). EM communities at unburned sites had the highest average evenness values of all site types. NMDS of fungal community dissimilarity (Bray-Curtis) showed that samples clustered primarily according to burn status and depth (*Figure 5*). EM communities were more like each other at burned than at unburned sites. Soil saprotrophs were more similar at unburned sites than at burned sites.

**Figure 5.**
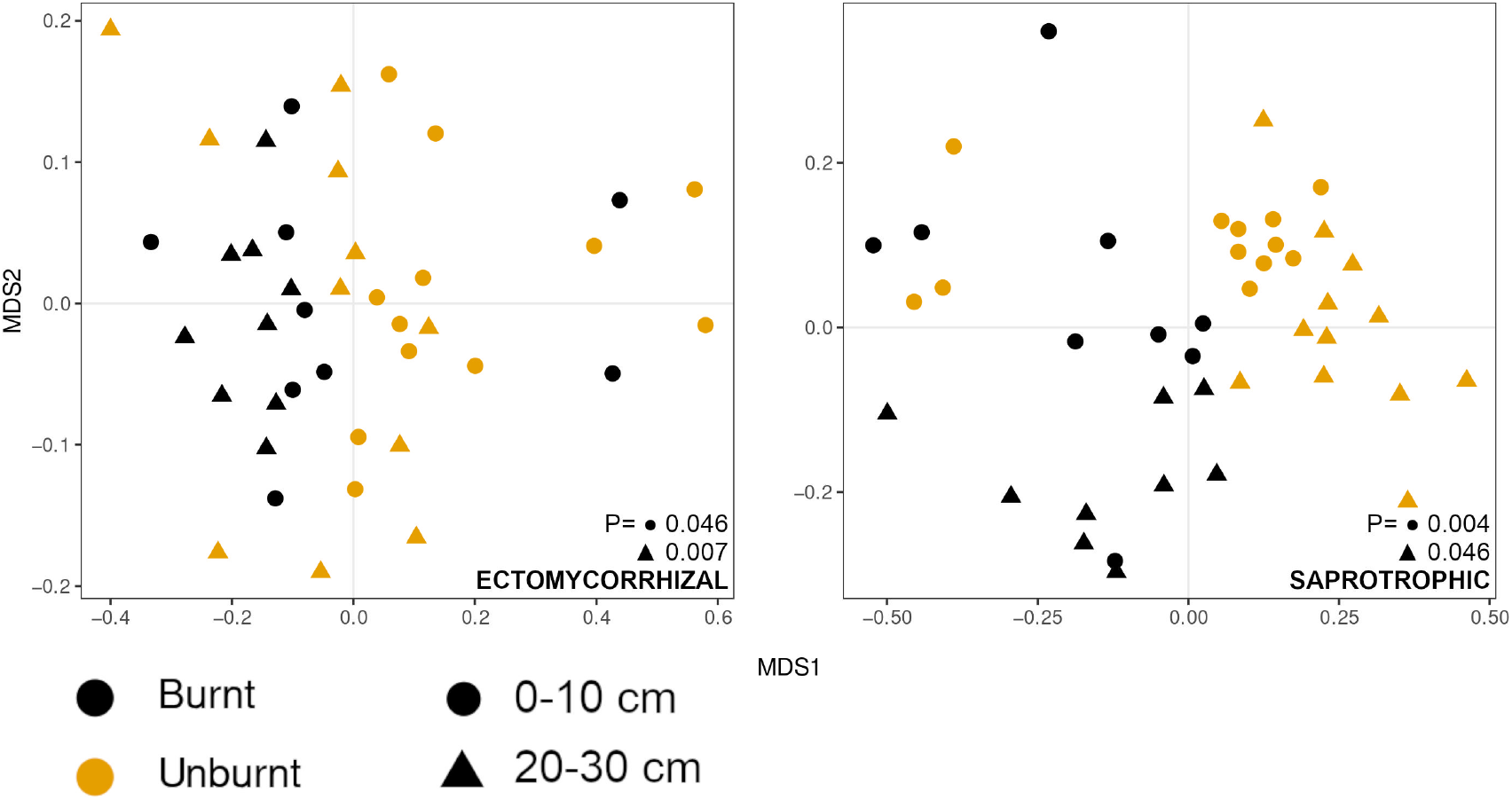
Non-metric multidimensional scaling showing differences in fungal community composition between burned and unburned sites at two sampling depths for ectomycorrhizal and saprotrophic fungi. Matrix based on Bray-Curtis distances. *P* values refer to PERMANOVA test for significance in differences in taxonomic community structure between burned and unburned sites at each sampling depth. (*Singlecolumn fitting image. Color to be used in print and on the Web*.)

### 3.5. Abundance of specific fungal taxa at sites with constrasting nutrient profiles

Relationships were detected between the abundance of some EM OTUs and levels of soil nutrients (*Figure 6*). *Russula*^3^, Pezizaceae, Pyronemataceae and Agaricales^5, 6^ were present in higher abundance in soil with elevated levels of N, P, K and Ca. *Entoloma* and *Tomentella* were associated with elevated pH, whereas *Sebacina, Chloridium* and Thelephoraceae 1 were associated with lower pH. Saprotrophic taxa associated with elevated N, P, K and Ca included *Bionectria, Leohumicola* and *Archaeorhizomyces* 6. Lower levels of these nutrients were associated with *Geminibasidium* 1, 2 and 3 and Apiosporaceae. Increased pH was associated with the saprotrophic taxa *Clavaria, Idriella* and Phallaceae, and lower pH with *Hygrocybe, Sakaguchia, Chaetosphaeria* and Thelephoraceae 2.

**Figure 6.**
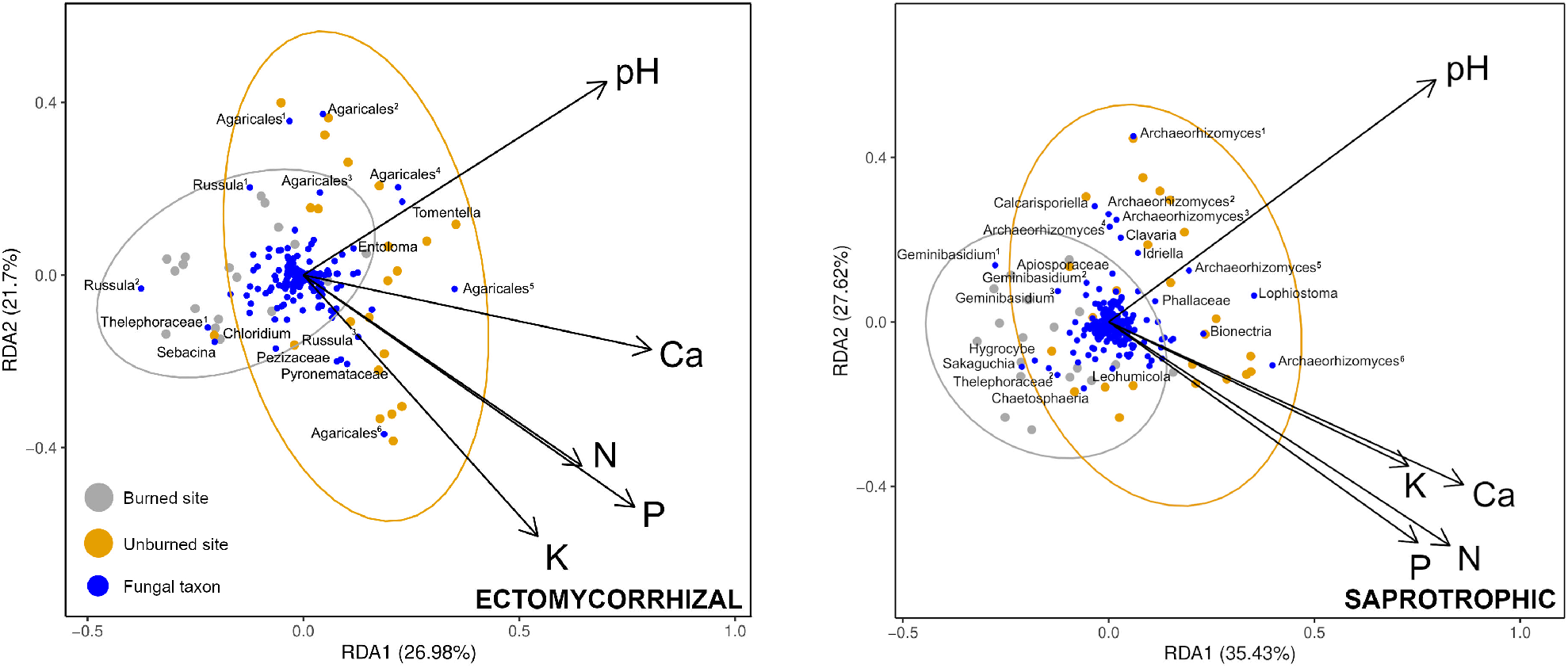
Redundancy analysis (RDA) summarising variation in response variables (fungal community composition and individual taxa) and explanatory variables (soil chemical/physical properties) at burned and unburned sites. Dark blue dots represent individual fungal operational taxonomic units; grey and yellow dots summarise differences in fungal community composition at burned and unburned sites. Taxon and site dots appearing closer to arrow tips were associated with higher levels of that soil chemical/physical variable, whereas taxon and site dots appearing opposite to an arrow were associated with lower levels. An angle of 90 degrees indicates little or no correlation. Based on Hellinger-transformed OTU tables to give lower weights to rare taxa. (*2-column fitting image. Color to be used in print and on the Web*.)

### 3.6. Abundance of specific fungal taxa at sites with constrasting fire histories

We assigned ecological guild and trophic mode for 165 EM and 654 saprotrophic soil taxa from 1,696 assigned OTUs with FUNGuild v1.1 (Nguyen et al., 2016). The basidiomycete taxa *Russula, Hemileccinium, Lactifluus, Amanita, Mycosymbioces* and Thelephoraceae were dominant at unburned sites and sites burned prior to 2015 (*Figure 7*). Ascomycota were present in increased abundance and taxonomic diversity, including *Meliniomyces* and the truffle taxon *Ruhlandiella*, at sites burned immediately prior to sampling in 2017. Species richness of EM fungi decreased linearly from the time of last fire, from 72 OTUs at unburned sites to 12 at sites burned in 2014. The exceptions to this trend were sites that were most recently burned, which had 58 EM OTUs.

**Figure 7.**
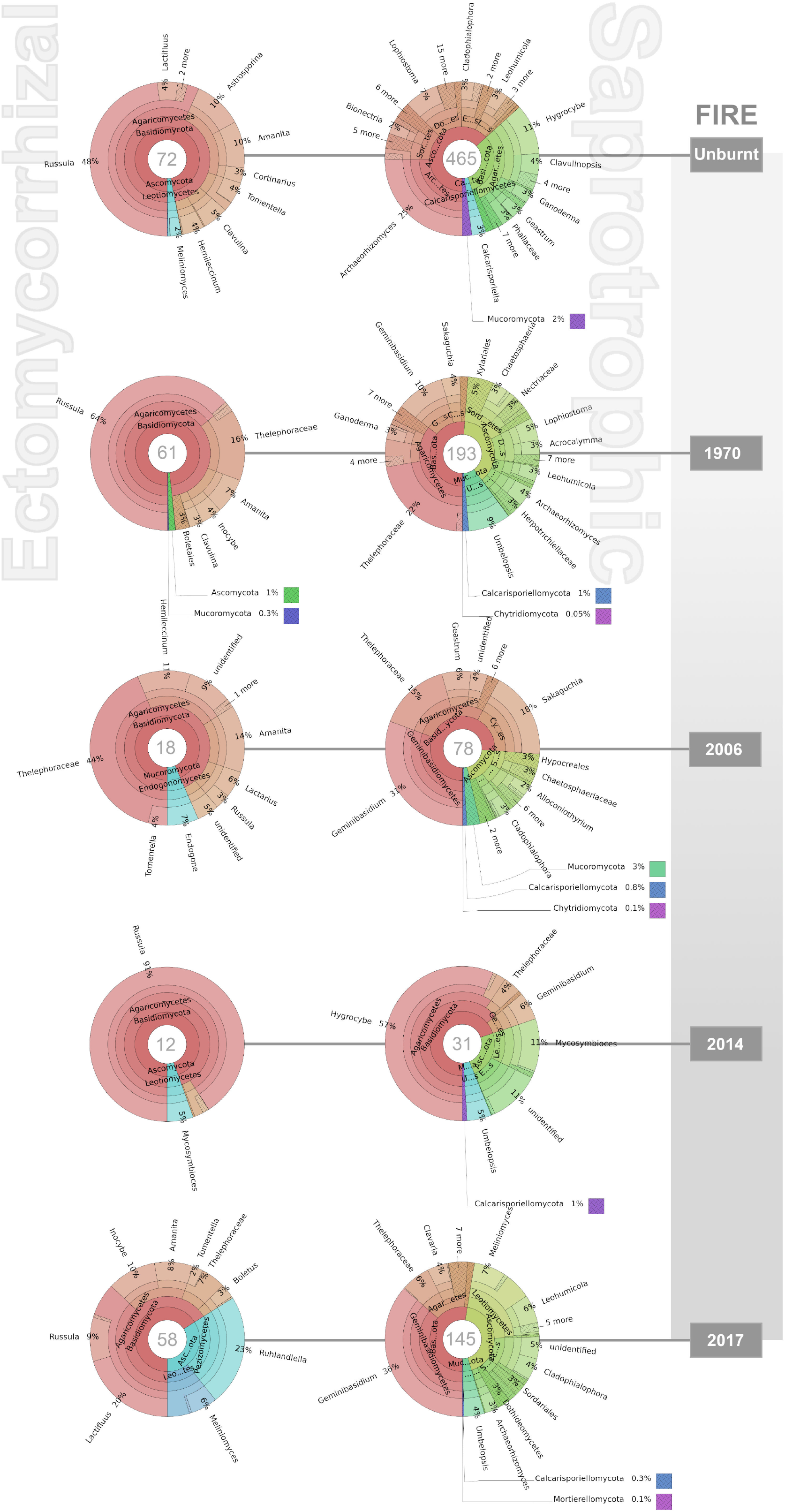
(previous page) Taxonomic composition and abundance of ectomycorrhizal (EM) and saprotrophic soil fungi at sites with different fire histories. Numbers inside circles in grey represent numbers of OTUs detected (species richness). Species richness of EM fungi decreased linearly from the time of last fire, from 72 OTUs at unburned sites to 12 at sites burned in 2014. (*2-column fitting image. Color to be used in print and on the Web*.)

Species richness was at least twofold higher for saprotrophs over EM at all sites, and up to 6 times higher at unburned sites (*Figure 7*). Unburned sites were dominated by *Archaeorhizomyces* (Ascomycota) and *Hygrocybe* (Basidiomycota). Dominant saprotrophic taxa at burned sites included *Geminibasidium, Hygrocybe*, Thelephoraceae (Basidiomycota) and *Umbelopsis* and *Mycosymbioces* (Ascomycota). Decreased species richness from the time of last fire was also evident for saprotrophs, from 465 OTUs at unburned sites to 31 at sites burned in 2014, with 145 OTUs at sites that were most recently burned.

## 4. Discussion

Saprotrophic and EM fungi showed major community-level differences at sites with contrasting fire histories, as did nutrient levels and composition of the dominant vegetation. EM composition was most different at burnt sites that contained mycorrhizal hosts. The changes to EM composition were largely unassociated with differences in soil chemistry. By contrast, the community composition, diversity and evenness of saprotrophic fungi was dissimilar between burned and unburned sites, and this was correlated with reduced soil nutrient levels. We also detected fire-associated differences in the composition of saprotrophic fungi in litter at shallow soil horizons.

### 4.1. Dominant plant communities influence ectomycorrhizal fungi

EM fungi are important tree symbionts in many forest ecosystems. The taxonomic community composition and abundance of EM fungi differed between burned and unburned forest soils, which was largely due to burned sites that contained the mycorrhizal hosts *Corymbia intermedia, Acacia flavescens and Acacia midgleyi*. In general, EM communities were less diverse and more even at unburned sites, which indicated a late-successional community structure dominated by a small number of taxa including *Russula* and *Amanita*. Successional shifts, in which EM community composition progresses from post-fire tree stand initiation to canopy closure, are well-documented (LeDuc et al., 2013; Longo et al., 2011).

The more recently a site had been burned, the lower the species richness of EM fungi, which was congruent with reports from temperate forests in the Northern Hemisphere (Kipfer et al., 2011; LeDuc et al., 2013; Rincón et al., 2014). Higher EM species richness at sites burned immediately prior to sampling contrasts with other studies that showed immediate negative effects of fire on EM diversity. One explanation may be that ascomycete EM taxa were more diverse and abundant at burned sites relative to unburned sites. If fire had occurred the year prior to sampling, *Ruhlandiella* (hypogeous fungi, or truffles), which are dispersed primarily by mycophagous mammals (Claridge, 2002; Dundas et al., 2018), were the most abundant of this pyrophilic group. *Ruhlandiella* are known to fruit abundantly after bushfires (Kraisitudomsook et al., 2019; Warcup, 1990). Reduction of undergrowth by fire also has the potential to increase mammalian access to the soil, which increases foraging and dispersal activity. Increased activity of pyrophilic taxa such as *Ruhlandiella* and those dispersed by mycophagous mammals may explain the short-term, postfire increase in EM diversity.

### 4.2. Saprotrophic fungal diversity was associated with nutrient differences between burned and unburned sites

This study showed that soil chemistry differed between burned and unburned sites, with structurally different saprotrophic communities associated with contrasting nutrient levels in the soil, and contrasting composition of the litter layer associated with different plant community composition. Saprotrophic fungi were more diverse and less even at unburned sites, where levels of soil nutrients were higher. Unlike EM fungi, saprotroph diversity was correlated with levels of all soil nutrients measured, including a weak yet measurable correlation with pH. Strong positive correlations between diversity of saprotrophic fungi and the soil quality indicators N, P and NH_4_ have been reported (Chen et al., 2021), as well as increases in saprotrophic biomass and diversity in response to experimental addition of N to soils (Clocchiatti et al., 2020). In a global study, Ca was found to be the strongest predictor of soil fungal diversity (Tedersoo et al., 2014). In this study in the AMT, we found Ca, NO_3_, P and K were strongly correlated with saprotroph diversity.

Soil saprotrophs showed marked changes in species richness that were congruent with a site’s fire history. In recently-burned areas, the species richness of soil saprotrophs was almost double that measured at most other burned sites. Fungal saprotrophs were dominated by *Geminibasidium* (Basidiomycota) and *Meliniomyces* (Ascomycota) in recently burned areas. We found higher species richness and Shannon’s diversity of saprotrophic over EM fungi regardless of a site’s fire history. Salo et al. (2019) described an increase in saprotrophic fungal diversity immediately after fire, with saprotrophic succession in soil more rapid than in wood. In Australian Mountain Ash forests, distinctive communities of soil fungi appeared in the year after fire disturbance, followed by much longer seral phases dominated by non-pyrophilic species (McMullan-Fisher et al., 2002). As outlined by Verma and Jayakumar (2012), low-intensity fires in 2017 may have increased the amount of organic material available, leading to a rise in saprotrophic diversity.

The dominant vegetation type was associated with saprotrophic communities in the litter soil horizon, which indicated that certain types of organic litter may favour some fungal taxa over others. Wu et al. (2011) found that leaf type was one of the main drivers of fungal community biomass and composition. Lunghini et al. (2013) reported higher fungal diversity in mixed litter than in monospecific litter. We propose that fire-induced alterations to plant community composition lead to compositional changes in the litter layer, which in turn select for particular communities of saprotrophic fungi.

### 4.3. Soil chemistry differed between burned and unburned sites

Chemical/physical properties differed between burned and unburned sites and between the litter and mineral soil layers, with NO_3_, P, Ca and pH generally higher in unburned than at burned sites. This is congruent with current knowledge of fire-nutrient dynamics in tropical forests, where soil nutrients are depleted by recurrent fires (Bowman, 2009a). Conversely, occasional fires can cause a short-term increase in nutrient availability at shallow soil horizons via combustion of litter and soil organic matter. Low wind and high sub-canopy moisture generate fires of lower intensity in AMT forests than in savannas (Cochrane, 2003; Verma and Jayakumar, 2012). Vegetation changes can influence levels of soil nutrients, especially N (Evans et al., 2001; Zhou et al., 2018). We found a greater net association of burning with soil chemical/physical properties, with no discernible patterns attributable to the different vegetation types studied. This suggests that alteration of nutrient profiles by fire has been direct, most likely through volatilization of litter and soil organic matter (Verma et al., 2019), rather than by indirect alterations to plant community composition.

We detected a strong correlation between levels of soil NH_4_ and the diversity of EM fungi. There was no correlation between EM diversity and other soil chemical/physical properties measured in this study. We propose that host availability and fire have a greater influence over rainforest EM community composition than levels of soil NO_3_, P, K, C, Ca and pH.

### 4.4. Different fungi at different depths

Vertical partitioning of fungi as observed in this study applies broadly to EM-dominated soil ecosystems in tropical and boreal zones (McGuire et al., 2013). Dominant vegetation was most strongly associated with EM communities in deep soil. The correlation of EM community structure with vegetation was weaker in the shallow litter layer, which is expected given the affiliation of EM fungi with tree roots. Primary notophyll rainforest in other areas of North Queensland has higher root biomass and root length compared to secondary forest (Hopkins et al., 1996). Deep soil horizons in unburned forests provide greater opportunity for EM colonisation of compatible hosts.

## 5. Conclusions

Our results supported the hypothesis that tropical soil fungi are impacted by burning, which was associated with changes in abundance and phylogenetic structure of EM and saproptrophic communities. Communities of EM fungi were structurally different at burned and unburned sites, which was associated with differences in the dominant vegetation, in particular with dominant EM taxa tolerant of frequent burning. Truffle-like taxa that are reliant on mycophagous mammals were more abundant at recently burned sites. In general, EM fungi at unburned sites had a late-successional community structure dominated by a small number of taxa. At burned sites EM diversity was higher and less even than at unburned sites. The diversity of saprotrophic fungi was associated with reduced soil chemical/physical levels at burned sites. In the litter layer, the community composition of saprotrophs was correlated with changes in vegetation type.

Globally between 2007 and 2017, carbon sinks provided by terrestrial ecosystems removed an estimated 32 % of anthropogenic CO2 emissions from the atmosphere (Le Quéré et al., 2018). Of these terrestrial sinks, tropical forests are some the largest due to their rapid growth ! (Keenan and Williams, 2018). Large savanna-dominated areas of Australia’s tropical north could, if protected from burning, support tropical forestry for carbon sequestration (K. Cook, pers.comm.), which may become a serious option for Australia as states begin to commit to net zero emissions (NSWDPIA, 2020). Any assessment of native tropical tree species for their utility in carbon forestry should consider their mycorrhizal symbionts and their tractability for the production of inoculum. Australia’s tropical fungi have the potential to serve as a major biological resource over the approaching decades.

## CRediT authorship contribution statement

**Jed Calvert:** Conceptualization, Methodology, Formal analysis, Investigation, Writing - Original Draft, Visualization, Project administration. **Alistair McTaggart:** Conceptualization, Methodology, Writing - Review & Editing, Supervision. **Lília C. Carvalhais:** Methodology, Formal analysis, Writing - Review & Editing. **André Drenth:** Writing - Review & Editing, Supervision. **Roger Shivas:** Writing - Review & Editing, Supervision.

## Research data for this article

The dataset and metadata supporting this article is available in the BioPlatforms Australia project’s data portal under the sample accessions 42144–42185 (https://ccgapps.com.au/bpa-metadata/base), doi:10.4227/71/ 561c9bc670099.

## Acknowledgements

The authors would like to acknowledge the Maxim Foundation for financial support for the project. We would also like to express our gratitude to Sofi Houghton for her design of the graphical abstract.

